# Reproductive capacity of *Varroa destructor* in four different honey bee subspecies

**DOI:** 10.1101/621946

**Authors:** Richard Odemer

## Abstract

Varroa tolerance as a consequence of host immunity may contribute substantially to reduce worldwide colony declines. Therefore, special breeding programs were established and varroa surviving populations investigated to understand mechanisms behind this adaptation. Here we studied the reproductive capacity in the three most common subspecies of the European honey bee (Carnica, Mellifera, Ligustica) and the F2 generation of a varroa surviving population, to identify if managed host populations possibly have adapted over time already. Both, singly infested drone and worker brood were assessed to determine fertility and fecundity of varroa foundresses in their respective group. We found neither parameter to be significantly different within the four subspecies, demonstrating that no adaptations have occurred in terms of the reproductive success of *Varroa destructor*. In all groups mother mites reproduce equally successful and are potentially able to cause detrimental damage to their host when not being treated sufficiently. The data further suggests that a population once varroa tolerant does not necessarily inherit this trait to following generations after the F1, which could be of particular interest when selecting populations for resistance breeding. Reasons and consequences are discussed.

**HIGHLIGHTS:**

- Varroa reproduction has been investigated in four different honey bee subspecies
- No differences could be observed, neither in drone nor in worker brood
- In managed bee populations, host immunity against *V. destructor* has not been established

**GRAPHICAL ABSTRACT:** Illustrations after Gullan & Cranston 2014

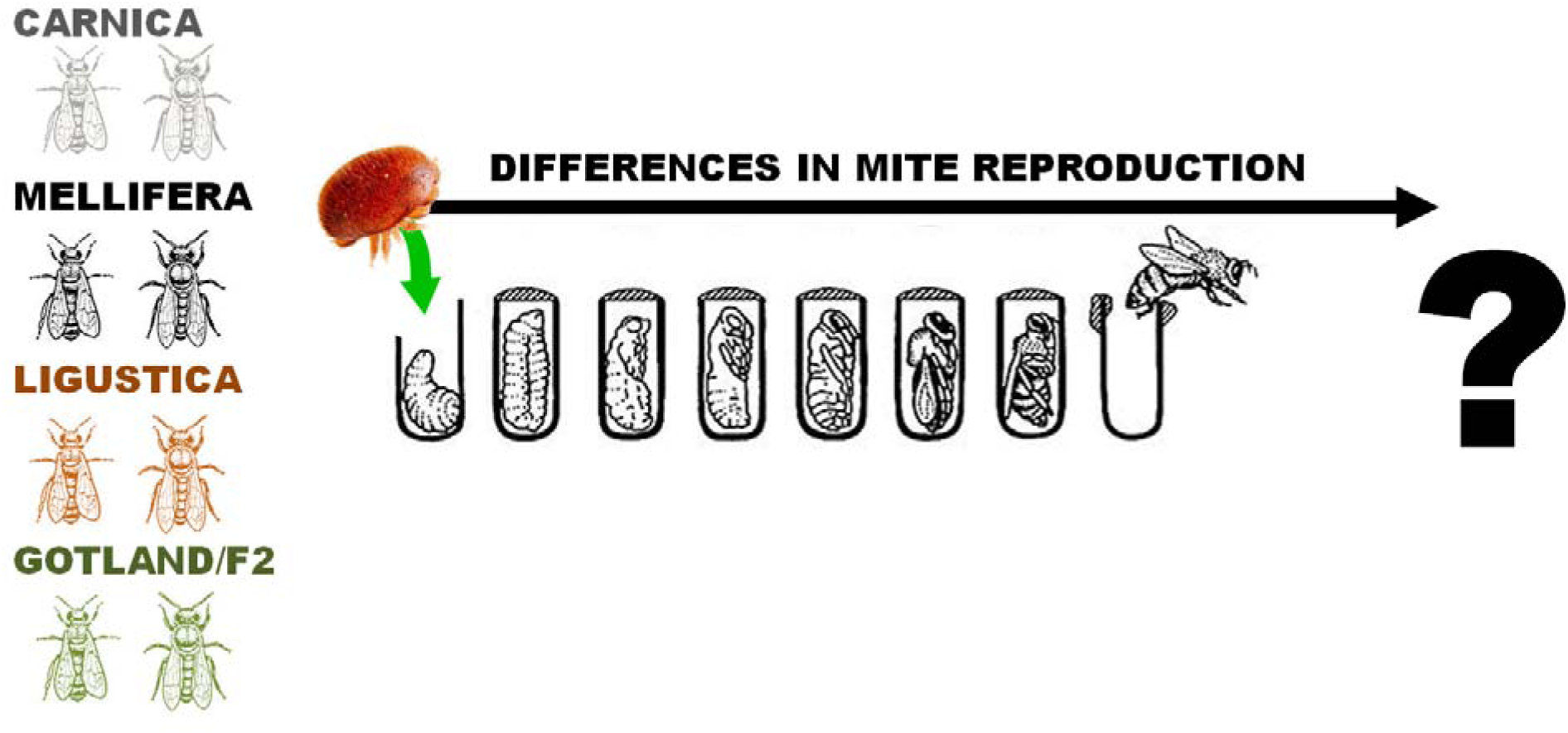

## 1 INTRODUCTION

Varroosis is known to be the most serious threat for European honey bees across the globe (Rosenkranz et al. 2010). A key for the mite’s success lies in their ability to perfectly adapt to host conditions, including the reproduction in worker brood. Even though reproductive capacity of *V. destructor* seems equally high in both, drone and worker brood, a distinctive amount of mites fail to reproduce even though they are not infertile (de Ruiter 1987). The conditions however, under which mite foundresses remain “temporary sterile” cannot yet be explained (Garrido & Rosenkranz 2003) but is discussed to be a host-specific tolerance trait against the mite (Rosenkranz & Engels 1994). Host stages in which mites are able to reproduce vary between drone and worker brood and reproduction is only possible within a narrow time frame, indicating a particularly sensitive process (Frey et al. 2013). Interestingly, Xie et al. (2016) revealed that mother mites are able to choose nurse bees over foragers and newly emerged bees as their optimal host in the phoretic phase, not only enabling them to quickly infest new brood cells (Donzé et al. 1998), but also providing the best possible nutritional conditions to produce a larger amount of progeny. Again, this demonstrates how highly adapted the parasite is.

Reports from surviving populations have increased over the last decade, suggesting a rapid host adaptation more or less simultaneously (Oddie et al. 2018). Besides a specific varroa mite targeted hygienic behavior (VSH = varroa sensitive hygiene) (Panziera et al. 2017), reduced mite reproduction is considered to be one key advantage for colony survival by means of natural selection (Locke et al. 2012). Almost exclusively, such traits have been investigated and documented for resistant honey bee populations (Locke 2016a) but have probably been neglected for more common subspecies. To close this knowledge gap and ascertain both, fertility and fecundity as a consequence of the reproductive capacity of *V. destructor*, we have compared the three most common subspecies of the European honey bee (Carnica, Mellifera, Ligustica) and the F2 generation of a varroa surviving population descending from the “Bond Project” on Gotland (Fries et al. 2006), to identify if managed host populations possibly have adapted over time already despite systematic control measures.

## 2 MATERIALS & METHODS

### Bee colonies and subspecies

A total of 22 honey bee colonies (*Apis mellifera* L.) were investigated during summer season from May to August. We focused on subspecies originating in Europe such as the Carniolan bee *A. m. carnica* (n=5), the European dark bee *A. m. mellifera* (n=7), the Italian bee *A. m. ligustica* (n=5) and a F2 generation of mite surviving bees from the “Bond Project” descending from the Swedish island of Gotland “Gotland/F2” (n=5). To provide a sufficient amount of drone pupa, one to two drone-frames were placed at the edge of the brood nest of each colony. All experimental hives were kept and maintained at our local apiary near the Apicultural State Institute in Stuttgart, Germany.

### Mite reproduction

The reproductive capacity of the foundress mite is specified as success to generate at least one viable daughter before the host pupa hatches (fertility). In contrast, mother mites that lay no or only a single egg, have no males or are delayed in egg-laying respective to host-development will fail to produce viable offspring for the following mite generations. Further, the number of progeny per mite (fecundity) serves as measure for a possible host adaptation representing a reduced reproductive capacity in terms of an increased survivability of the colony.

To increase comparability of our results, all experiments were performed according to the methods described in Locke & Fries (2011). In brief, worker and drone pupae in stage Pd and older, but before eclosion, were examined (see Fig. 1). At least 30 cells per colony were carefully investigated where possible and mite infestation was documented. Only cells with a single foundress were considered, cell content and mites attached to the pupa were accurately removed and subsequently observed under a stereo-microscope (Zeiss Stemi 2000-CS).

**Fig. 1.**
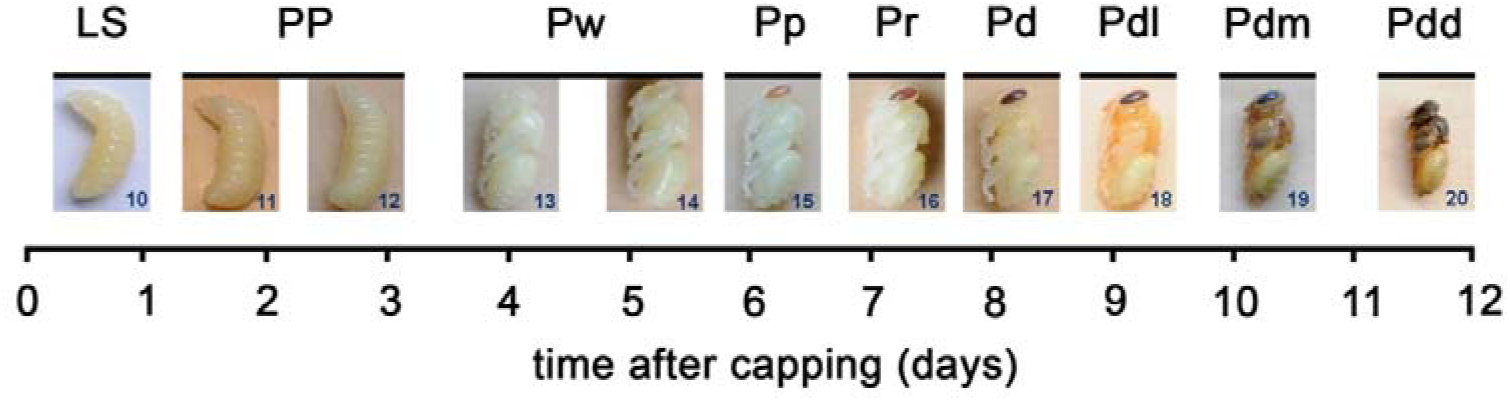
Classification of pupal stages respective to ontogenetic worker development (after Rembold et al. 1980, graphically modified after Wang et al. 2015). Abbreviations: LS = 5^th^ larval instar after sealing; PP = prepupa; P = pupa (w = white eyes; p = pink eyes; r = red eyes; d = dark brown eyes; dl = dark brown eyes, light pigmented thorax; dm = dark brown eyes, medium colored thorax; dd = dark brown eyes, dark thorax).

### Data evaluation

Mite reproduction and fecundity data were first tested for variance homogeneity and normal distribution with Levene’s and Shapiro-Wilk test and verified for both datasets, respectively. A generalized linear model was applied to both sets followed by a comparison of the least-squares means and a *P* value adjustment (tukey method). For all tests RStudio (R Core Team, 2018) and significance level of α=0.05 was used.

## 3 RESULTS

Different parameters of varroa mite reproduction in four different honey bee subspecies are presented in Table 1. A total of n = 3104 drone and n = 2526 worker brood cells were evaluated. We did not find significant differences for the overall reproductive capacity (fertility) in the four groups. Neither in worker brood (df = 10: *F* = 2.26; *P* = 0.144) nor in drone brood (df = 15: *F* = 2.51; *P* = 0.098). A similar outcome was observed for the average number of offspring per foundress (fecundity). Both, progeny found in worker brood (df = 10: *F* = 2.84; *P* = 0.092) and in drone brood (df = 10: *F* = 2.32; *P* = 0.873) were at the same level.

Due to an increased infestation rate which resulted in a high ratio of multiply infested cells in the drone brood of all four subspecies, it was not possible to evaluate drone pupa in stage Pd and older as previously described. To compare fecundity regardless these circumstances, we had to consider earlier developmental stages beginning already at Pw (Fig. 1) providing a sufficient amount of singly infested cells. This is why the average number of offspring is relatively low when compared to worker brood.

**Tab. 1.** Comparison of the reproductive capacity (fertility and fecundity) of mother mites produced in singly infested drone and worker brood cells

|  | Carnica | Mellifera | Ligustica | Gotland/F2 |  |
| --- | --- | --- | --- | --- | --- |
| **Drones** |  |  |  |  |  |
| No. of cells (n) | 68 | 179 | 51<sup>b</sup> | 141 |  |
| Fertility | 79% | 83% | 59% | 79% | *ns* |
| Fecundity (± SE)<sup>a</sup> | 2.7 (± 0.5) | 2.7 (± 0.3) | 2.2 (± 0.6) | 2.6 (± 0.2) | *ns* |
| **Workers** |  |  |  |  |  |
| No. of cells (n) | 90 | 91 | 120 | 120 |  |
| Fertility | 82% | 89% | 96% | 78% | *ns* |
| Fecundity (± SE) | 3.3 (± 0.3) | 3.4 (± 0.3) | 4.1 (± 0.4) | 3.3 (± 0.2) | *ns* |

a: earlier developmental stages beginning already at Pw had to be considered for the drone brood
b: not representative, due to the low amount of singly infested cells (10 cells per colony on average)
ns: not significant ( $P > 0.05$ )

For the number of cells in Ligustica drone brood it needs to be mentioned that due to the late re-queening of experimental colonies (mid July) it was not possible to obtain a sufficient amount of singly infested cells. Hence, we only used 10 cells per colony on average, this should be considered when interpreting the results.

## 4 DISCUSSION

Here, we studied the reproductive capacity of three commonly managed honey bee subspecies and the F2 generation of a varroa surviving population originated from the “Bond Project” (Fries et al. 2006). When compared to former data, the fertility of varroa foundresses in worker brood did not change significantly during the past three decades and has levelled off between 80 to 90 % (Thrybom & Fries 1991; Corrêa-Marques et al. 2003; Alattal et al. 2006; Locke et al. 2012; Alattal et al. 2017). This trend is corroborated by our data and most likely similar for drone brood.

Drone frames that we have investigated here were highly infested already in early summer, not least because some colonies remained untreated in the former season at our experimental apiary but also because the mite’s preference to infest drone cells is approximately eight times higher when compared to worker brood (Fuchs 1990; Santillán-Galicia et al. 2012). In addition, the time frame which is attractive to enter cells for infestation is approximately twice as long in drone brood (Calderone et al. 2002), being one reason for this preference. Under these circumstances it was not surprising that we found many multiply infested drone cells and it became a challenge to locate cells containing only one foundress for our evaluation. Ligustica queens arrived after summer solstice very late in the season and besides that, a very high mite infestation in drones was the reason that we were not able to collect a sufficient amount of singly infested cells.

Moreover, our data confirms that there is no large selection pressure favoring reduced mite reproduction in both, drones and workers, at least not under intensively managed conditions. For the three common subspecies this is not remarkable as host adaptations are most often reported as a means of natural selection (Seeley 2007; Locke et al. 2012; Oddie et al. 2017). For the F2 generation of the surviving population from Gotland however, we had expected a different outcome. The Gotland bees have developed an apparent reduced mite reproductive success trait that is either inheritable from paternal, maternal or both sides in the F1 generation (Locke 2016b). Our results provide evidence that this trait seems to fade out by further generational change, once more making the colonies susceptible to *Varroosis*.

Although we did not find significant differences in the fertility and fecundity of varroa females between surviving F2 and common honey bee subspecies, we are still convinced that the varroa reproductive capacity represents a crucial and probably the only parameter for the future selection of varroa resistance on the individual level. One reason is that we confirmed that about 85 % of the “temporary sterile mites” were again fertile if re-introduced into freshly sealed brood cells (Weller 2008). Hence, the occurrence of “temporary sterile mites” seems to be rather a trait of the host than a trait of the parasite and, therefore, offers possibilities for selection.

## 5 CONCLUSION

Frequent reports have shown that apart from the most common managed honey bee subspecies there are populations demonstrating increased mite susceptibility and great variance in mite reproductive capacity (de Guzman et al. 2008; Locke 2016a; Nganso et al. 2018). This reflects an encouraging potential to establish varroa resistance in European *A. mellifera* populations (Büchler et al. 2010). However, resistance mechanisms are complex which is why further research is necessary to understand host-adaptation and mite reproduction in greater detail.

